# Recapitulation of clinical features in a patient-derived xenograft mouse model of VEXAS Syndrome

**DOI:** 10.1101/2024.08.27.609907

**Authors:** Liliana Arede, Rita Coutinho, Tiago L Duarte, Marta Lopes, Rui V Simões, Susana Silva, Pedro Baptista, José Manuel Lopes, Raquel Faria, Delfim Duarte

## Abstract

Patient-derived xenografts (PDX) are valuable tools to model human disease and test new therapies. Here, we report, for the first time, a PDX model recapitulating the unique characteristics of a VEXAS patient (unique patient number UPN1).

## Report

In late 2020, a new syndrome affecting predominantly adult males with myelodysplasia and autoinflammatory symptoms was described^1^. The new entity termed VEXAS (Vacuoles, E1 enzyme, X-linked, Autoinflammatory, Somatic) Syndrome results from somatic mutations in UBA1 (Ubiquitin-like modifier-activating enzyme 1) on the X chromosome within hematopoietic stem and progenitor cells. Initially, three variants of this mutation were described in exon 3: p.Met41Leu, p.Met41Thr and p.Met41Val. The *UBA1* gene has two isoforms, *UBA1a* (nuclear) and *UBA1b* (cytoplasmatic). More recently, *UBA1* splice-site mutations have also been described as pathogenic. Described mutations lead to a new splicing variant *UBA1c* and reduced levels of *UBA1b*, which is associated with decreased ubiquitination activity and results in a pro-inflammatory cellular state^2^.

Clinical features of VEXAS include weight loss (54.5%), skin lesions (83.6%), chondritis (36.2%), pulmonary involvement/infiltrates (40.5-49.1%), and ocular involvement (40.5%) ^3^. Hematologic changes include macrocytic anemia and thrombocytopenia. A diagnosis of myelodysplastic syndrome and/or monoclonal gammopathy of unknown significance is established in up to 50% of patients^3^. It is unclear whether UBA1 mutated clones directly cause the autoinflammatory changes or indirectly result from a pro-inflammatory milieu. VEXAS patients are generally treated with corticosteroids and immunosuppressive drugs such as tocilizumab (Interleukin-6 inhibitor) and Janus Kinase (JAK), mainly ruxolitinib. However, these approaches are only partially effective, and VEXAS has high mortality rates. There is an urgent need to understand VEXAS pathophysiology and for novel treatments.

Patient UPN1 was diagnosed at the age of 68 years old with recurrent polychondritis, bilateral episcleritis, leukocytoclastic vasculitis, recurrent rash, trilineage myelodysplasia, deep vein thrombosis, IgG/k monoclonal gammopathy, and elevated erythrocyte sedimentation rate (ESR) and C reactive protein, later confirmed as VEXAS Syndrome, with the presence of the *UBA1* p.Met41Leu variant.

During bone marrow (BM) assessment, 7×10^6^ primary BM mononuclear cells were obtained and intravenously transplanted into a sublethal irradiated immunodeficient NSG-SGM3 (NSGS) recipient mouse. The animal was monitored, and we could detect peripheral blood engraftment (0.72%), anemia and thrombocytopenia (**Supplementary Figure 1A, B**) from 3 months post-transplantation. From 7 months post-transplantation, the animal showed health deterioration with progressive weight loss, ptosis, and reduced mobility. At eight months, the animal was euthanized, and we were able to detect human hematopoietic cell engraftment (hCD45+) in the BM (0.5%) and spleen (9%), as evaluated by flow cytometry (**Figure 1A**). The NSGS recipient (VEXAS-PDX) presented macrocytic anemia, thrombocytopenia (**Figure 1B**), pale BM, hypercoagulability, and splenomegaly (**Figure 1C**). We performed tissue histological analysis confirming the presence of hCD45 cells in the spleen and lung (**Figure 1D**). In the lung, there was destruction of alveolar spaces and infiltration by immune cells, including polymorphonuclear leukocytes (**Figure 1E, white arrows**), resembling alveolitis, not observed in the control (**Figure 1E**). Moreover, the VEXAS-PDX presented defects in the cartilage compatible with chondritis, as suggested by magnetic resonance imaging (MRI), namely, cartilage denudation and apparent degenerative changes of the joint, compared to the control animal (**Figure 1F**). The xenograft also reliably recapitulated the auto-inflammatory profile with increased human IL-6 production (**Figure 1G**). Interestingly, we also detected an increase in mouse TNF and mouse IL-6, suggesting the contribution of a bystander effect (**Figure 1H**). The main features of the VEXAS-PDX and the corresponding UPN1 are described in **Table 1**.

**Table 1.**

|  | **UPN1** | **VEXAS-PDX** |
| --- | --- | --- |
| **UBA1 mutation** | p.Met41Leu (c.121A>C) | p.Met41Leu (c.121A>C) |
| **Sex (M/F)** | M | M |
| **Elevated inflammation** | Yes | Yes |
| **Macrocytic anemia** | Yes | Yes |
| **Thrombocytopenia** | Yes | Yes |
| **Pulmonary involvement** | Yes | Yes |
| **Chondritis** | Yes | Yes |
| **Deep vein thrombosis** | Yes | Hypercoagulability |
| **Skin involvement** | Yes | Not detected |
| **Ocular involvement** | Yes | Ptosis |
| **Vasculitis** | Yes | Not assessed |

**Figure 1.**
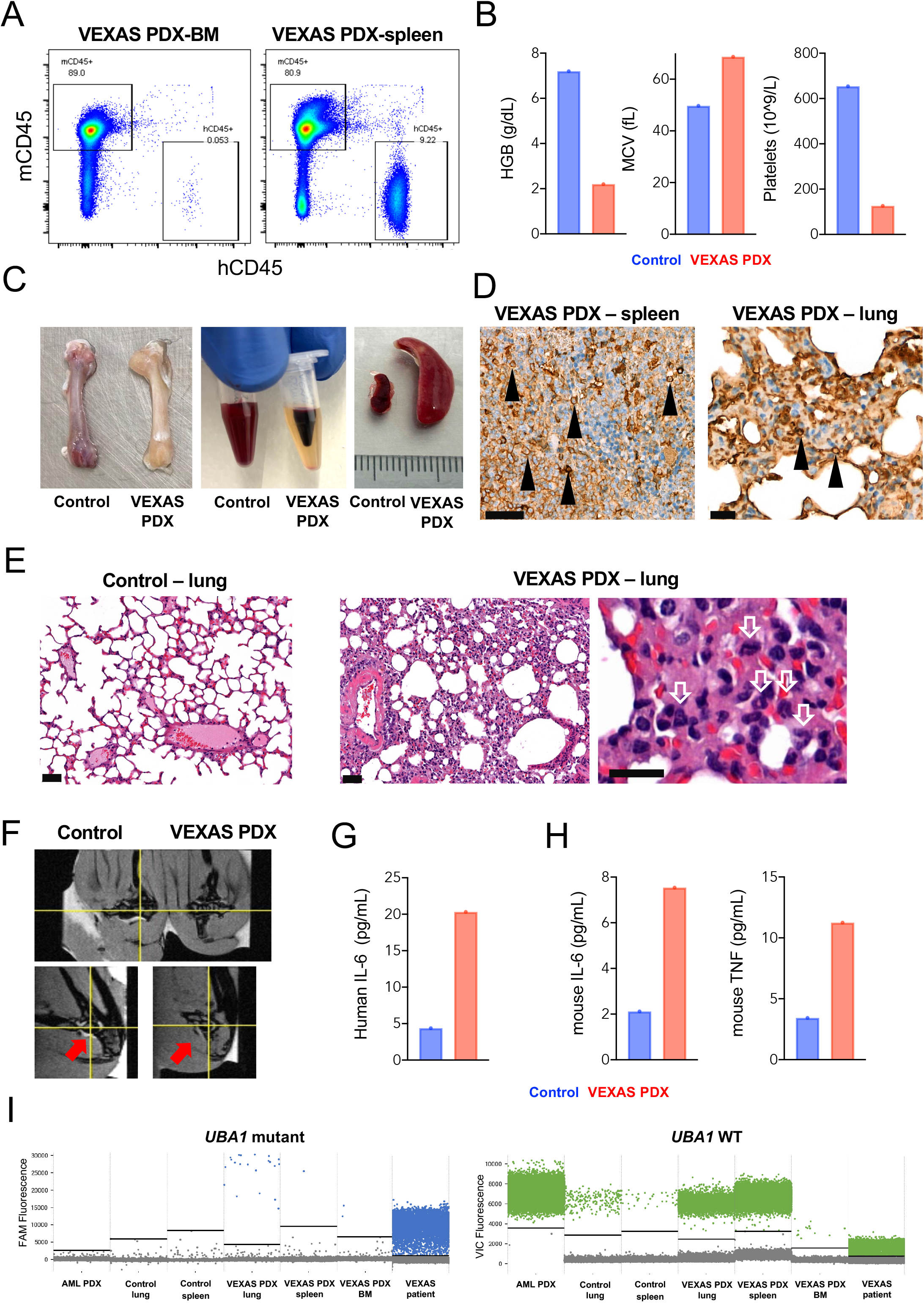
Development of hematological, immunophenotypic, and morphological features of VEXAS in a PDX mouse model. **(A)** Engraftment of human CD45 (hCD4+) cells in the bone marrow (BM) and spleen. **(B)** Assessment of the parameters hemoglobin (HGB), mean corpuscular volume (MCV) and platelet counts. **(C)** At 8 months, VEXAS PDX presented with severe anemia (pale bones), hypercoagulability and splenomegaly. **(D)** Immunohistochemistry revealed infiltration of the spleen and lung with hCD45+ cells. Scale bars: 60*μ*m (left) and 20*μ*m (right) **(E)** In contrast with the control (left), histological analysis of the VEXAS PDX lung (right) revealed alveolitis with the presence of polymorphonuclear cells. Scale bars: 60μm (left and middle) and 20μm (right) **(F)** *Ex vivo* MRI of the joints revealed loss of cartilage and signs of degeneration. **(G)** Human IL-6 and **(H)** mouse IL-6 and TNF were significantly increased in the serum of the VEXAS PDX. **(I)** Digital PCR using mutation-specific probes revealed the presence of UBA1-mutated cells in the VEXAS PDX (negative controls: AML PDX, control mouse; positive control: VEXAS patient).

Engrafted cells carried the genetic variant of the corresponding patient, as confirmed by digital PCR (**Figure 1I**), with an approximate Variant Allele Frequency (VAF) of 13.45% in the BM, 0.0047% in the spleen and 0.4992% in the lung. An additional negative control of an hAML PDX without the UBA1 variant was used.

We establish that transplantation of primary BM cells from a VEXAS patient into immunodeficient NSGS mice can recapitulate the clinical features of the syndrome. Our study is limited by the single mouse used but proves that UBA1-mutated cells can engraft and that the human phenotype can be partially recapitulated in mouse recipients. NSGS mice are severely immunodeficient transgenic mice with defective innate (low natural killer (NK) cells, macrophage activity, absent complement system) and adaptive (lack of B and T cells) immune functions^4^. The model was genetically modified to express the human growth factors SCF, GM-CSF, and IL-3, thus facilitating the engraftment of primary cells derived from patients^4^. These features make NSGS mice ideal recipients for patient-derived cells, and they have been used to model hematological diseases notoriously challenging to study, including chronic myelomonocytic leukemia (CMML)^5^. A recent study has shown that monocytes are essential for establishing an inflammatory environment in VEXAS^6^.

Interestingly, GM-CSF is crucial for the pro-inflammatory activation of monocytes. Whether VEXAS primary cells are susceptible to SCF, GM-CSF, or IL-3 should be explored in the future. In the present case, unselected MNCs were transplanted. Transplantation of unmutated cells may be necessary for the observed phenotype. Still, future studies should test pre-selection of CD34+ cells and intrafemural injections as strategies to increase the success of PDX establishment and reduce the lag time of clinical manifestations^5^.

In our model, we detected severe cytopenias and extramedullary hematopoiesis that likely reflect the loss of normal hematopoiesis in the animal. It would be interesting to use humanized mice, previously engrafted with healthy CD34+ cells and then transplanted with VEXAS samples, to explore how UBA1 mutated clones affect normal hematopoiesis. It should be noted that although we detected higher engraftment of hCD45 in the spleen, VAFs for UBA1 mutation in hCD45 populations were higher in the BM, and BM cellularity was low. These findings suggest that UBA1 mutated clones disrupt usual BM niches and that unmutated UBA1 hCD45 cells expand better in an extramedullary environment, namely the spleen.

Inflammation may be a crucial factor in impaired hematopoiesis^7^ observed in VEXAS. We detected high levels of human IL-6; these cells are intrinsically primed for cytokine production. Using the xenotransplantation setting, we measured mouse cytokines in a cytometric bead array (CBA) assay and detected elevated mouse IL-6 and TNF. This reveals that inflammation in VEXAS patients likely results from contributions of UBA1 mutated cells but also from cell extrinsic sources. We detected infiltration of affected tissues, such as the lung, by mutated hCD45 cells, suggesting that UBA1 mutated clones are direct effectors of tissue pathology, either causing damage per se or locally recruiting inflammatory cells from the host (e.g., neutrophils).

Another relevant point of our observations is the potential use of VEXAS PDX models as avatars for patients to predict clinical evolution. In the present case, patient UPN1 had no manifestations of lung involvement at the time of cell collection and PDX establishment. Indeed, our PDX model predicted alveolitis, which was later diagnosed in the patient.

Altogether, our model captures the unique immunophenotypic and morphological features of VEXAS. Our data reveal that the NSGS mouse can sustain the development of VEXAS-initiating cells *in vivo*, setting up a putative new pre-clinical model that may be useful to investigate disease pathophysiology and develop effective therapies.

## Acknowledgments

The authors thank the patient for his participation in this study. We also thank the veterinarian (Sofia Lamas) and the caretakers from the i3S Animal Facility; Maria Lázaro (b.IMAGE, i3S) for technical assistance; Mafalda Rocha (i3S Genomics Core Facility) and Rui Batista (Thermo Scientific) for assistance with digital PCR; Carlos Bilreiro (Champalimaud Foundation) for discussing the MRI images.

