## Supplemental Materials for "Recapitulation of clinical features in a patient-derived xenograft mouse model of VEXAS Syndrome"

**Supplementary Materials and Methods**

**Patient clinical characteristics**

After informed consent, the primary cells were obtained from a patient recruited at Centro Hospitalar Universitário de Santo António, Porto. All experiments conformed to the World Medical Association (WMA) Declaration of Helsinki principles. For patient clinical characteristics, see **Table 1**.

**Establishment of the patient-derived xenograft**

The patient-derived xenograft (PDX) was generated by intravenous injection of 7x10^6^ primary VEXAS cells in a NSGS (NOD.Cg-Prkdcscid Il2rgtm1Wjl/SzJ) 8 week-old male, previously conditioned with sub-lethal irradiation of (2.4 Gy) 3h before transplantation. Another sublethally irradiated sex and age-matched control was also used. Animals were housed in an individually ventilated cage with access to food and water *ad libitum*. VEXAS-like progression was monitored in the peripheral blood by flow cytometry using anti-mCD45 and anti-hCD45 antibodies. To minimize suffering, mice were euthanized upon VEXAS-recipient reaching a predefined humane endpoint. VEXAS cells were collected from spleen and bone marrow. Mice were treated in accordance with the European Union Directive 2010/63/EU. The procedures were previously approved by the Portuguese authority Direção-Geral da Alimentação e Veterinária (DGAV), as well as by the i3S animal ethics committee (DD_2019_15).

**Flow cytometry**

For mouse BM analysis, femurs and tibias were dissected and crushed in phosphate-buffered saline containing 2% fetal bovine serum. Splenocytes were obtained by mashing mouse spleens in the same buffer. After red blood cell (RBC) lysis (RBC lysis buffer 1×, Cat. No. 420302, BioLegend), BM and spleen cells were transferred through a 40-µm cell strainer into a 50-mL tube and centrifuged at 500g, 4°C for 5 minutes. Cells were stained for 15 minutes at RT with the anti-human CD45 antibody (Cat. No. 304012, BioLegend) and anti-mouse CD45 antibody (Cat. No. 103116, BioLegend) at 1:200 dilution. Samples were run on a flow cytometer (FACS Canto II, BD Biosciences) and data were analyzed with FlowJo (BD Biosciences).

**Hematological profiling**

Mouse blood cell parameters were evaluated on an HT5 Veterinary Hematology Analyzer (Heska) using 15-20 µL of peripheral blood or BM aspirates.

**Digital PCR**

According to the manufacturer's instructions, genomic DNA was extracted from mouse tissues using an NZY Tissue gDNA isolation kit (MB13502, NZYtech) and quantified on a Qubit 2.0 Fluorometer (Invitrogen). Detection of the UBA1 mutation p.Met41Leu (c.121A>C) in mouse tissues was performed on an Absolute Q digital PCR (analysis of 20,000 micro-chambers per reaction) system (Thermo Fisher Scientific) using a Taqman SNP Genotype assay (4331369). A FAM probe detected the mutation, whereas a VIC probe detected the WT allele. Data were analyzed with QuantStudio™ Absolute Q™ Digital PCR Software. VAFs$were calculated as follows: \%=\frac{FAM (copies/\mu L)}{VIC (copies/\mu L )+ FAM (copies/\mu L )} x 100$

**Cytokine analyses**

Human interleukin 6 (IL-6) levels in mouse serum were measured using the ELISA MAXTM Standard Set (Cat. No. 430501, BioLegend) following the manufacturer’s instructions. The absorbance was determined at 450 and 570 nm on a µQuant Universal Microplate Spectrophotometer (BioTek, MQX200). Levels of cytokines in mouse serum were determined using the Cytometric Bead Array (CBA) mouse inflammation kit (Cat. No. 552364, BD Biosciences) according to the manufacturer’s instructions. Data were acquired on a flow cytometer (FACS Canto II, BD Biosciences) and analyzed with FCAP Array software from BD Biosciences.

**Histologic analysis**

Mouse tissue samples were fixed in neutral formalin 10% and paraffin-embedded. Staining with hematoxylin and eosin (H&E) and the immunodetection of human CD45 (using monoclonal mouse anti-human CD45, clones PD7/26 + 2B11, DAKO cat. #M0701, at 1:600 dilution) were performed at the Department of Pathology, Centro Hospitalar Universitário São João.

***Ex vivo* MRI**

The right upper limbsfrom VEXAS PDX and CONTROL mice were excised following euthanasia and immediately frozen at 20ºC. The tissues were thawed 1 week later and mounted on a 20 mL syringe, pre-filled with Fluorinert. The sample was loaded on a 3 Tesla Bruker Maxwell scanner (Bruker BioSpin, Ettlingen, Germany), equipped with high-power gradients (900 mT/m) and a cryorefrigerated radiofrequency coil, for sensitivity enhancement. After standard MRI adjustments, 3D MRI data were acquired from the joint region: first, with a T1-weighted gradient-echo pulse sequence (FLASH: 50/4.5 ms TR/TE, 15º flip angle, 100 µm isotropic resolution, and 2.5 min acquisition time); then, with a T2-weighted balanced steady state free precession pulse sequence (trueFISP: 7/3.5 ms TR/TE, 30º flip angle, 80 µm isotropic resolution, and 2 min acquisition time; sequence repeated with 60 µm isotropic resolution and 2.5 min acquisition time). The raw data was then loaded in Matlab (R2023b, Natick MA, USA) and exported in Nifti format for further analysis by a Radiologist.
