## Supplementary figures and images for "Recapitulation of clinical features in a patient-derived xenograft mouse model of VEXAS Syndrome"

### Supplementary Figure

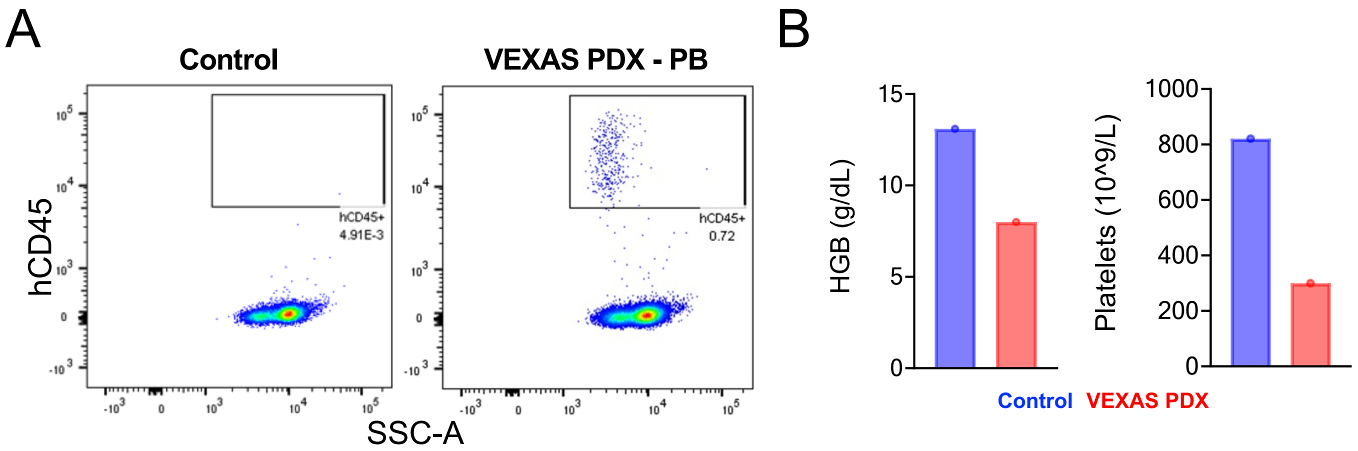
